# Pyrrole produced by *Pseudomonas aeruginosa* influences olfactory food choice of *Caenorhabditis elegans*

**DOI:** 10.1101/2022.01.27.477966

**Authors:** Deep Prakash, Ritika Siddiqui, Sreekanth H. Chalasani, Varsha Singh

## Abstract

Sense of smell can influence dietary choices in animals. So far, most of the research has focused on how animals respond to distinct odors when they are introduced individually. However, it remains unclear how animals evaluate foods that contain a bouquet of olfactory cues with contrasting effects. Here, we utilize *Caenorhabditis elegans* as a bacterivore to ask if odors produced by dietary bacteria can regulate worms’ food preferences. We show that the bacterium *Pseudomonas aeruginosa* produces a relatively small quantity of a new attractant for *C. elegans*. We identify the odor as a heterocyclic compound called pyrrole. We find that pyrrole contributes to the sensory decision-making of worms in diet preference assays. Using specific neuronal ablation lines and calcium response assays, we show that AWA odor sensory neurons of worms are necessary for sensing pyrrole. In all, we show that specific odors produced by bacteria can influence food choice behavior of animals.

## INTRODUCTION

Animals make specific dietary decisions to optimize caloric intake based on their needs ^[1]^. Food quality can be evaluated using specific sensory cues such as taste and smell. Pairing of food with specific smells is known to significantly alter food intake in mice ^[2]^. For bacterivores, such as *C. elegans*, several diet choices are available in their natural habitat of decaying organic matter ^[3-5]^. A well-developed chemosensory system including an odor sensory system enables worms to efficiently engage in food search behavior as well as avoid pathogens ^[6-9]^. *C. elegans* displays chemotaxis response to a wide range of volatile molecules including alcohols, ketones, amines, aldehydes, organic acids, aromatic and heterocyclic compounds ^[10]^. Many of these are products of bacterial secondary metabolism and encompass attractive cues such as diacetyl produced by lactic acid bacteria ^[11]^ as well as repulsive cues such as 2-nonanone. Some of the attractive odors are likely used as food cues by worms ^[12]^.

*C. elegans* habitat is composed of bacteria producing both attractive and repulsive odors. Often a single bacterium can produce contrasting olfactory signals. For example, *Serratia marcescens* causes an aversion response in worms but also produces attractive odors such as acetone and 2-butanone ^[9, 13]^. In the case of *P. aeruginosa*, 24h old lawn produces a repulsive odor, 1-undecene, that induces an aversion response in worms ^[14]^. However, worms prefer *P. aeruginosa* when presented with a choice between *Escherichia coli* and a 5h lawn of *P. aeruginosa* ^[6]^. This indicates that an attractive cue might also be produced by *P. aeruginosa*, at least in the young lawns of the bacterium. The identity of such an attractant remains unknown. It is also not clear how worms assess the quality of diet containing both attractive and repulsive cues.

Here, we studied the preference of *C. elegans* for *P. aeruginosa* over *E. coli*, to understand the contribution of various bacterial odors to the worm’s diet preference. We found that worms indeed prefer a young lawn of *P. aeruginosa* over that of *E. coli*. We analyzed the volatiles profile of young lawn of *P. aeruginosa* by gas chromatography mass spectrometry. By analyzing the worm’s response to each of the six individual odors present in the headspace of *P. aeruginosa*, we found small quantities of an attractive odor, pyrrole. We show that pyrrole is sensed by the AWA neurons. Adaptation experiments showed that pyrrole contributes to the preference of worms for young lawns of *P. aeruginosa*. Finally, we show that worms use AWA neurons to assess diet quality when both attractant and repellent, pyrrole and 1-undecene produced by *P. aeruginosa*, are present.

## RESULTS

### Pyrrole produced by *P. aeruginosa* is an attractant for *C. elegans*

*C. elegans* can distinguish between different bacteria and make appropriate diet choices. Given a choice between freshly plated (5h) laboratory food *E. coli* and pathogenic *P. aeruginosa*, worms show a slight preference for *P. aeruginosa* early on but learn to avoid it later ^[6]^. We found that wild-type N2 worms indeed displayed a modest but consistent preference for 8h young lawn of *P. aeruginosa* PA14 over *E. coli* OP50 in the diet preference assay (Figure 1A). To test whether this was due to olfactory cues produced by the dietary bacteria, we tested the requirement for AWA and AWC neurons of worms, two pairs of odor sensory neurons known to sense attractive odors ^[15, 16]^. Worms with a mutation in *odr-7*, a gene required for the specification of AWA neurons, did not display a preference for PA14, while worms carrying AWC ablation, AWC(-), showed a preference for PA14 over OP50 (Figure 1A). This suggested that AWA neurons of *C. elegans* contribute to worms’ preference towards the young lawn of PA14 over OP50.

**Figure 1.**
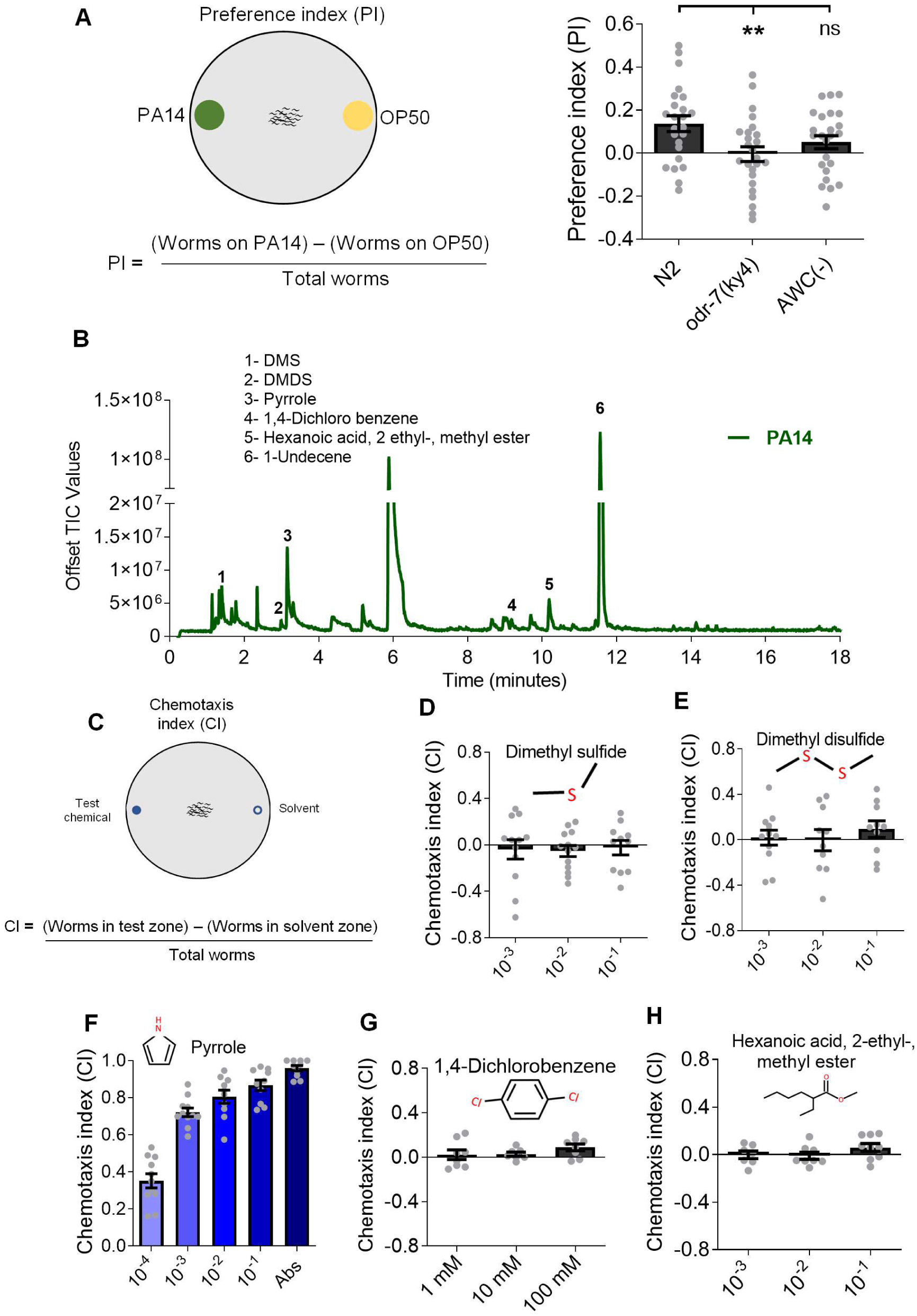
Pyrrole produced by *P. aeruginosa* serves as an attractant for C. elegans. (A) Schematic for the diet preference assay. Preference index (PI) of N2, *odr-7(ky4)*, and AWC(-) worms in diet preference assay between PA14 and OP50. (B) GC-MS profile of volatiles detected in the headspace of PA14 lawn. (C) Schematic for the chemotaxis assay. Chemotaxis indices of N2 worms to varying dilutions of (D) Dimethyl sulfide, (E) Dimethyl disulfide, (F) Pyrrole, (G) 1, 4-Dichlorobenzene (H) Hexanoic acid 2-ethyl-, methyl ester. ** P ≤ 0.001 as determined by a two-tailed unpaired *t*-test. Error bars indicate SEM. See supplementary figure S1 and movie S1.

To identify attractive odors produced by *P. aeruginosa*, we analyzed the volatiles present in the headspace of PA14 using an SPME fiber followed by gas chromatography mass spectrometry (GC-MS) as described before^[14]^. We found six volatiles in its headspace which were not present in the control, media only, headspace (Figure 1B and Figure S1). Of these, 1-undecene is an aversive cue that is sensed by AWB odor sensory neurons resulting in an aversion response of worms on an old lawn of *P. aeruginosa* ^[14]^. As expected, worms showed an aversion response to 1-undecene (Figure S2). To find attractants, we carefully examined the chemotaxis response of N2 worms to each of the remaining 5 volatiles at a two-fold range of concentrations (schematic in Figure 1C). Worms showed no response to dimethyl sulfide (DMS) or dimethyl disulfide (DMDS) (Figures 1D and 1E) consistent with a previous report ^[12]^. Worms also showed no response to 1,4-dichlorobenzene or hexanoic acid, 2-ethyl-, methyl ester (Figures 1G and 1H). However, pyrrole induced a strong dose-dependent chemotaxis response in worms (Figures 1F, also see Supplementary Movie S1).

How do worms assess bacteria such as *P. aeruginosa* producing both a repellent and an attractant, 1-undecene and pyrrole. We hypothesized that worms use ratiometric (attractant/repellent) odor code for chemotaxis. To test this hypothesis, we examined chemotaxis response of worms in different odor landscapes created using varying ratios of the two odors (schematic in Figure S3A). We observed that the presentation of absolute 1-undecene with an increasing concentration of pyrrole had a dramatic effect on the behavior of worms. The repulsion of worms to 1-undecene gradually altered to attraction with increasing levels of pyrrole. Thus, worms showed a range of behavior from chemorepulsion to chemoattraction with an increase in the attractant/repellent ratio (Figure S3B). To determine if worms respond to absolute levels of the odors or to their ratios, we presented the repellent at 3-fold range of concentrations and paired each with an increasing concentration of the attractant. In all cases, we were able to see a switch from chemorepulsion to chemoattraction (Figures S3B-E). This suggested that it is not the absolute amount but the ratio of the two contrasting cues that determines worms’ chemotaxis response (Figure S3F). Adding additional odors, DMDS, DMS, and 1,4-Dichlorobenzene, did not alter the ratio-tracking ability of N2 worms (data not shown). Altogether, our findings indicate that worms can detect precise ratios of attractant to repellent to arrive at a chemosensory decision in a complex odor-rich environment. We propose that this ratio is an odor-based code for diet quality.

### AWA odor sensory neurons control the chemotaxis response of worms to pyrrole odor

To test whether pyrrole is active as an odor, we performed a chemotaxis assay in a tripartite plate where pyrrole was partitioned from worms using a polystyrene barrier and it was only available as a volatile. Even under these conditions pyrrole induced a strong chemotaxis response in N2 worms (Figure 2A). Next, we tested whether worms’ odor sensory mechanisms are necessary for sensing pyrrole odor. To begin with, we examined the chemotaxis response of N2 and *odr-3* mutant worms to pyrrole. ODR-3 is a G protein alpha subunit active in *C. elegans* odor sensory neurons ^[17]^. We found that *odr-3(n2150)* mutants did not respond to pyrrole at any of the tested concentrations (Figures 2B, also see Supplementary Movie S2). This suggested that odor sensory neurons are necessary for sensing pyrrole. Further, to determine the identity of neurons necessary for sensing pyrrole, we used mutants or ablation lines for odor sensory neurons AWA, AWB, and AWC. A mutation in *lim-4*, necessary for the development of functional AWB neurons ^[18]^, or AWC ablation strain had no impact on worms’ chemotaxis response to pyrrole. This indicated that pyrrole sensing is independent of the sensory functions of AWB or AWC neurons (Figure 2C). An AWA ablation strain as well as *odr-7 (ky4)* mutant, both lacking functional AWA neurons, failed to respond to pyrrole (Figure 2C). This indicated that AWA odor sensory neurons are essential for responding to pyrrole. Diacetyl, a *bonafide* food cue and an odor produced by lactic acid bacteria, is detected via G protein-coupled receptor ODR-10, expressed in AWA sensory neurons ^[11, 19]^. We tested the chemotaxis response of the *odr-10(ky32)* mutant to pyrrole and found it to be comparable to that of N2 worms (Figure 2C) indicating that ODR-10 GPCR does not detect pyrrole. We also compared the dose-response of N2 worms to an increasing concentration of diacetyl and pyrrole independently and found the chemotaxis indices (CIs) to be comparable (Figure 2D). Altogether, our results suggested that pyrrole is a strong attractant and is sensed by AWA odor sensory neurons of *C. elegans*.

**Figure 2.**
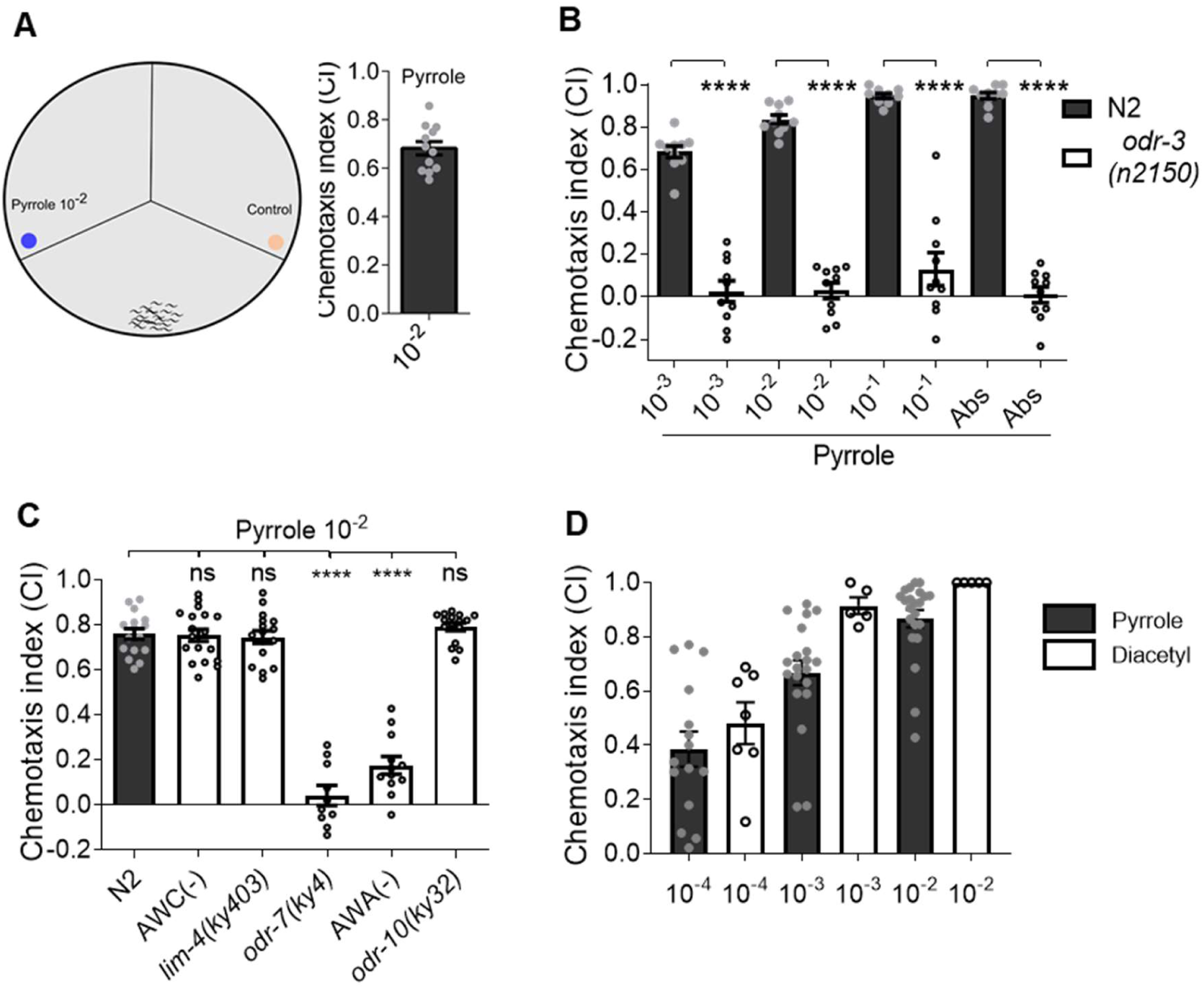
AWA odor sensory neurons control chemotaxis response to pyrrole. (A) Chemotaxis response of N2 worms towards pyrrole odor at 10^−2^ dilution. (B) Chemotaxis response of N2 and *odr-3(n2150)* worms to various concentrations of pyrrole. n ≥ 3 assays. N ≥10 assay plates with 30-50 worms in each plate. **** P ≤ 0.0001 as determined by two-tailed unpaired *t*-test. Error bars indicate SEM. (C) Chemotaxis of N2, AWC(-), *lim-4(ky403), odr-7(ky4)*, AWA(-), and *odr-10(ky32)* worms to pyrrole at 10^−2^ dilution. n ≥ 3 assays. N ≥10 assay plates with 30-50 worms in each plate. ns (not significant) P > 0.05, **** P ≤ 0.0001 as determined by one-way ANOVA, followed by Dunnett’s multiple comparison test. (D) Chemotaxis response of N2 worms to increasing concentrations of pyrrole and diacetyl. See movie S2.

### The odor of pyrrole activates calcium signaling in AWA odor sensory neurons

To confirm the role of AWA odor sensory neurons in sensing pyrrole, we examined the odor-evoked calcium response of AWA neurons ^[20]^. In worms expressing genetically encoded calcium sensor, GCaMP, in the AWA neurons, we observed a dose-dependent stimulus-response within 1-2s of the presentation of pyrrole at the nose of the worm (Figures 3, A-D). This is consistent with the requirement of AWA neurons for chemotaxis response to pyrrole (Figure 2C). We did not observe calcium response, to pyrrole, in AWC_on_ or AWC_off_ neurons (Figures S4, A-C), again consistent with their dispensability in chemotaxis assays. We observed a slight response in AWB neurons upon stimulus withdrawal, in some worms (Figures S4A and S4D). In all, these results confirmed that AWA neurons are not only necessary for the chemotaxis response to pyrrole, but they are also the primary sensing neurons for this odor.

**Figure 3:**
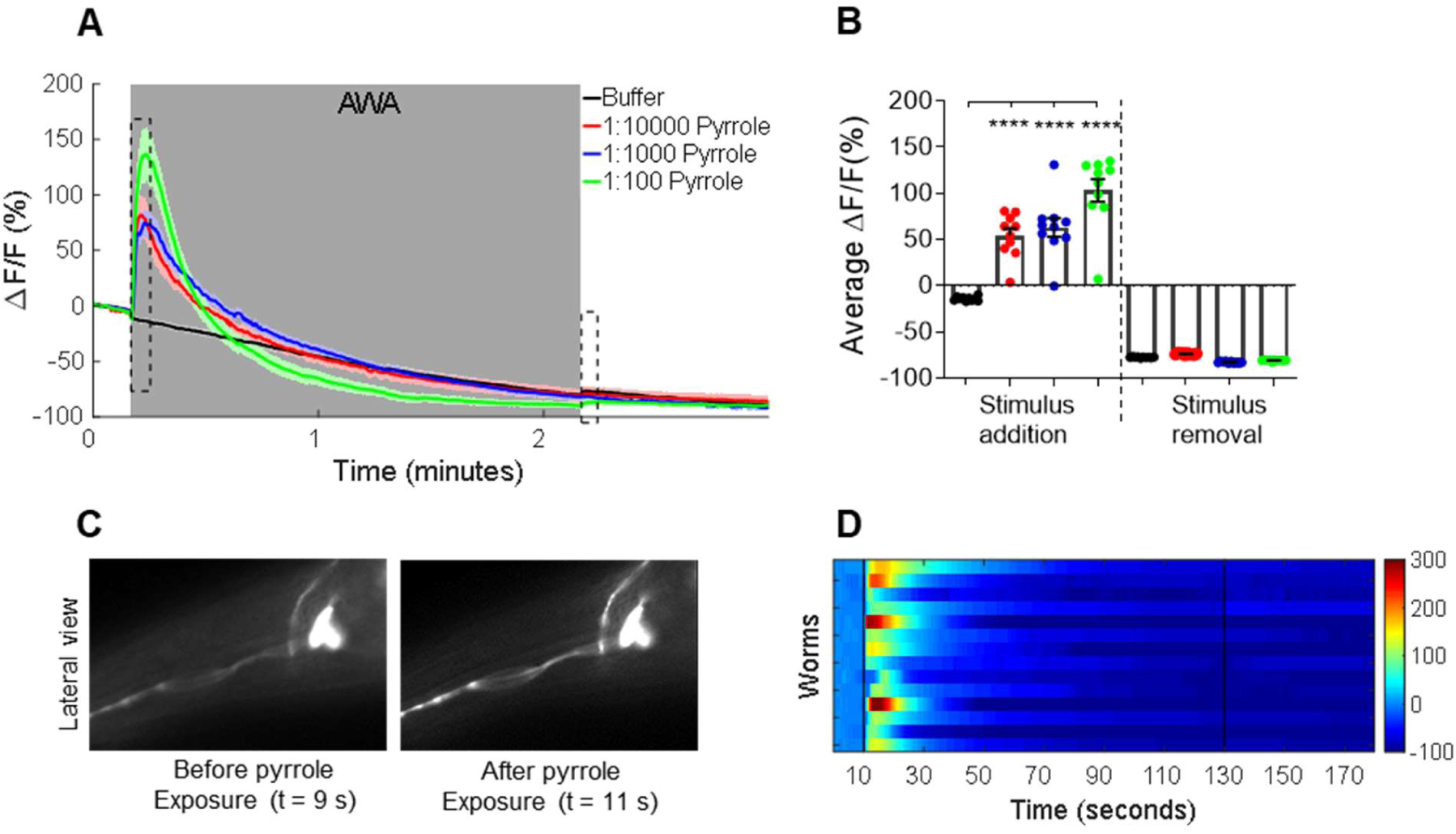
AWA odorsensory neurons are activated by the odor of pyrrole. (A) Average calcium responses of AWA::GCaMP2.2b worms to pyrrole, recorded for 180s, under stimulus between 11s-130s window shown in grey. (B) Average percentage change in ΔF/F for 10s time window after stimulus addition and withdrawal (dashed boxes in A), each data point represents average data of ≥ 13 worms. **** P ≤ 0.0001 as determined by one-way ANOVA, followed by Dunnett’s multiple comparison test. Error bars indicate SEM. (C) GCaMP2.2b fluorescence in AWA neuron before (9^th^s) and just after pyrrole exposure (11^th^s). (D) Heat map of calcium response in AWA::GCaMP2.2b worms. Each row represents an individual worm recorded for 180s under 10^−2^ dilution of pyrrole, in 11s-130s window. See supplementary figure S2.

Since AWA odor sensory neurons are primary sensing neurons for pyrrole, we asked whether these neurons are also needed to respond to attractant/repellent ratio. We tested the chemotaxis of N2 and *odr-7(ky4)* worms against a mixture of varying ratios of pyrrole and 1-undecene. As shown in Figure S5, while N2 worms were able to track the increasing ratio of pyrrole to 1-undecene, *odr-7* mutants failed to do so. This indicated that *C. elegans* requires AWA odor sensory neurons to modify its behavior in a complex, odor-rich environment. Altogether, our experiments suggested that *C. elegans* uses odor sensory neurons to detect precise ratios of the attractant to the repellent, likely mimicking alterations in the odors produced by bacteria during its growth.

### Pyrrole influences the dietary choice of worms for young lawn of PA14

To understand the relevance of *P. aeruginosa*-produced pyrrole for worms, we first asked if pyrrole can have an impact on the diet preference of worms. To do this, we took advantage of a well-established odor adaptation assay. Prolonged exposure of *C. elegans* to an odorant is known to induce adaptation leading to loss or reduction in the worm’s response to the same odorant in subsequent exposures ^[21]^. We used a modified adaptation assay ^[12]^, where we exposed worms to pyrrole for 90 minutes followed by testing their chemotaxis to pyrrole compared to naïve worms (Figure 4A). Worms adapted to pyrrole showed diminished chemotaxis response to pyrrole compared to naïve worms (Figure 4C) but displayed normal chemotaxis to diacetyl and 2-nonanone reflecting the specificity of adaptation (Figure S6). Next, we tested both naïve and pyrrole-adapted worms in a diet preference assay. We found that pyrrole-adapted worms lost the preference for 8h lawn of PA14 over OP50 compared to naïve worms (Figure 4B). These experiments suggested that pyrrole produced by *P. aeruginosa* contributes to the preference for the young lawn of PA14 displayed by worms in a diet choice assay.

**Figure 4.**
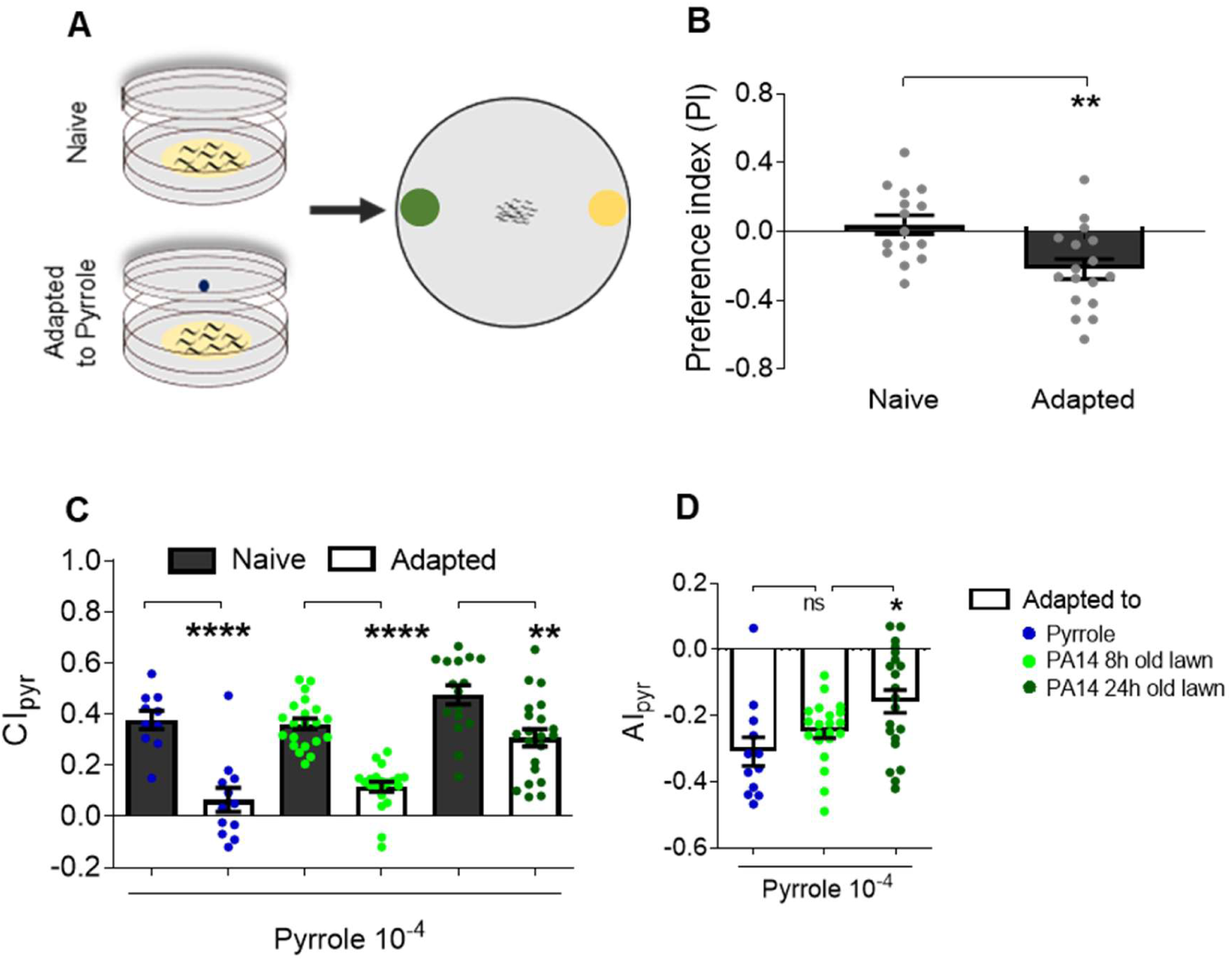
Pyrrole influences the dietary choice of worms for young lawn of PA14. (A) Schematic of adaptation assay with pyrrole followed by diet preference assay. (B) Diet preference index of naïve worms and worms adapted to pyrrole. (C) Chemotaxis response of naïve and adapted worms to pyrrole. Adaptations conditions are indicated in F. (D) Adaptation index for the data shown in E. *Adaptation Index (AI) = (Chemotaxis Index of Adapted worms) – (Chemotaxis Index of naïve worms)*. ns (not significant) P > 0.05, * P ≤ 0.05, ** P ≤ 0.01, **** P ≤ 0.0001 as determined by two-tailed unpaired *t*-test. Error bars indicate SEM.

We have earlier shown that *C. elegans* displays a significant preference for the young lawn over the older lawn of *P. aeruginosa* in an ODR-3 dependent manner ^[14]^. We hypothesized that a young lawn of PA14 (8h) has a higher level of pyrrole than in the older lawn (24h). Based on this, we predicted that adaptation with odors from 8h lawn of PA14 will dampen chemotaxis response to attractant pyrrole. To investigate age-dependent changes in the level of pyrrole, we performed adaptation of worms using odors either from a young lawn (8h) or from an older lawn (24h) of PA14. As controls, we also performed adaptations of worms with pyrrole alone. As expected, adaptation with odors from 8h lawn was as effective as an adaptation with pyrrole itself in dampening CI of worms while adaptation with odors from 24h lawn of PA14 had a much smaller effect on CI of adapted worms compared to naïve worms. We also calculated adaptation index to pyrrole (AI) for worms adapted to pyrrole, to odors of 8h lawn of PA14, or odors of 24h lawn of PA14. The AI of the first two groups were comparable while both were significantly different from the AI for worms adapted to odors from the older (24h) lawn of PA14 (Figure 4D). The decline in the level of pyrrole in the older lawn, inferred from adaptation experiments, is consistent with a previous report of decline in the level of pyrrole in aging *Pseudomonas* culture ^[22]^. Collectively, the adaptation experiments suggested that level of pyrrole is higher in the younger lawn of PA14 consistent with worms’ diet preference for PA14.

Overall, our results show that worms’ dietary preference for *P. aeruginosa* is driven by bacterial pyrrole sensed by AWA odor sensory neurons of worms (Figure 5).

**Figure 5.**
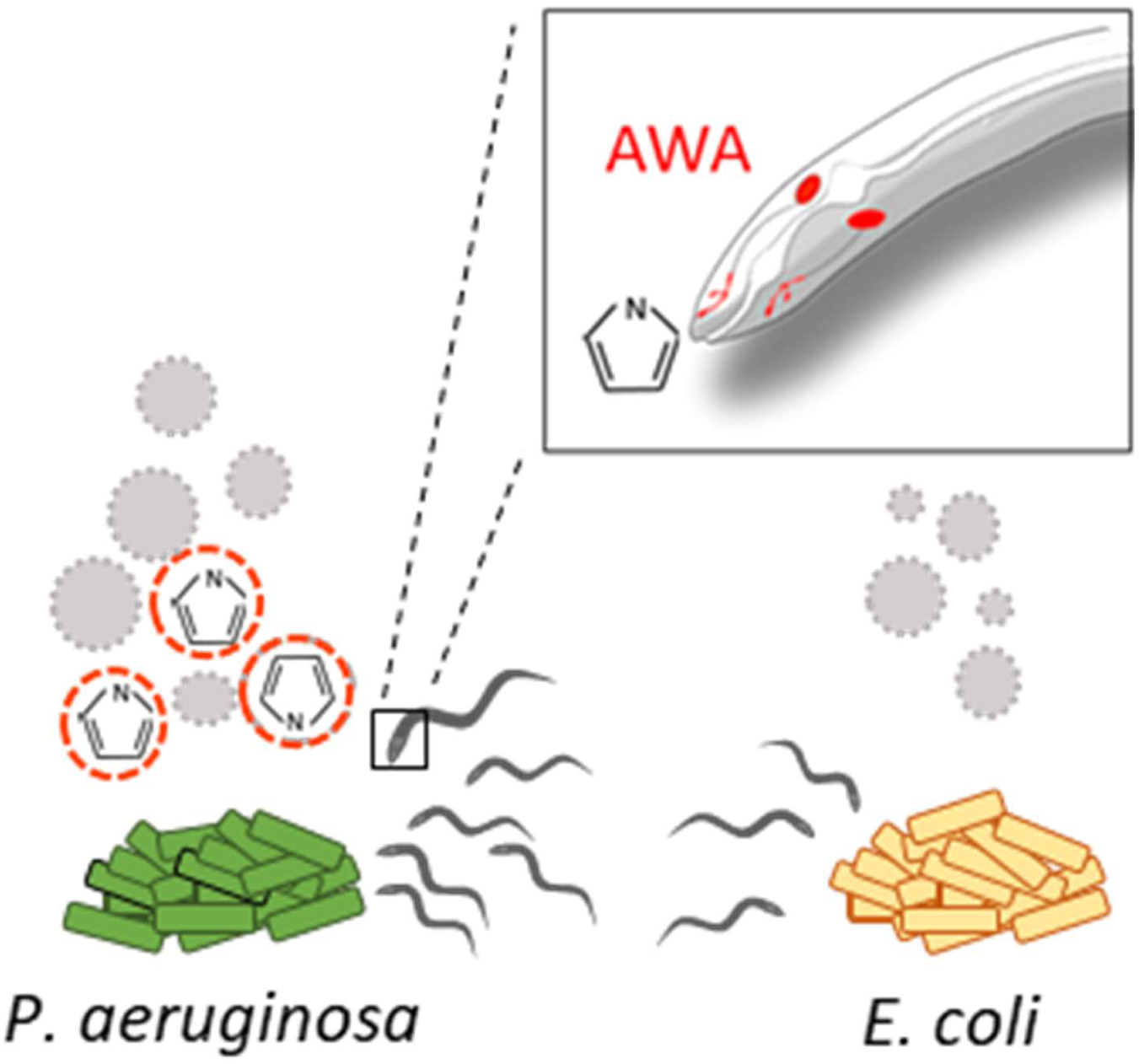
Pyrrole influences the dietary choice of worms for young lawn of *P. aeruginosa* over *E. coli* via the engagement of AWA odor sensory neurons of *C. elegans*.

## DISCUSSION

In this study, we have demonstrated the presence of odor tracking mechanisms in *C. elegans* which allows them to make appropriate diet choices in a complex, odor-rich environment. We show that pathogenic *P. aeruginosa* produces an attractant, pyrrole, in young lawns which makes them attractive to worms via the action of AWA odor sensory neurons. By changing the ratio of the attractant to the repellent, we could faithfully change *C. elegans* behavior from chemoattraction to chemorepulsion. This is an example of a robust chemosensory mechanism in animals enabling them to have better foraging capacity in nature where nutrient sources are limiting.

It is interesting to find an attractive odor, pyrrole, in a pathogenic bacterium *P. aeruginosa*. At least two other pathogenic bacteria are known to produce attractive volatiles to entice *C. elegans. S. marcescens* produces 2-butanone and acetone ^[12]^, while *Bacillus nematocida* B16 produces 2-heptanone ^[23]^. What is even more noteworthy is that the levels of volatiles are dynamic and vary with the age of bacteria. For example, pyrrole attains maximum levels during the early phase of *P. aeruginosa* growth and then declines ^[22]^. This is consistent with our adaptation experiments, in which we discovered that young *P. aeruginosa* lawns contained more attractant (pyrrole) while the older, aversion-inducing lawns had less of it. What is the relevance of pyrrole for bacteria and worms? One possibility is that *P. aeruginosa* uses pyrrole as a lure to attract *C. elegans*. Alternatively, the presence of pyrrole, a nitrogen-containing heterocyclic compound, in a young lawn is likely to be reflective of a rapidly growing (log phase) population of *P. aeruginosa*. Thus, pyrrole may be an indicator of good food for worms. High pyrrole/1-undecene might indicate bacteria in the log phase while low pyrrole/1-undecene might indicate bacteria in saturation phase where not only the bacterium is starving but quorum sensing is achieved, and virulence factors are produced ^[24]^. The food quality is likely coded in the nervous system of worms as the precise ratio of pyrrole/1-undecene for *P. aeruginosa*. The presence of *Pseudomonas* in the native microbiome of *C. elegans* as well as the presence of pyrrole and 1-undecene uniquely in the headspace of *P. aeruginosa* further supports the possibility of a *Pseudomonas* specific odor code. Although pathogenic, *Pseudomonas* species have indeed been found in the intestine of native *C. elegans* ^[4, 25]^ suggesting that worms do consume them.

What is the advantage of having an adaptable diet preference behavior? It is conceivable that odor-coded food preference is an evolved trait in worms in the face of seasonal and ephemeral foods. When food is limiting, the capacity to choose a palatable (low repellent, high attractant) diet over starvation could be a highly desirable trait to have. Indeed, food limitations have led to the evolution of several adaptable traits in organisms such as dauer formation in *C. elegans*.

Recent studies in *Drosophila melanogaster* and *C. elegans* have suggested that the presentation of two contrasting stimuli simultaneously can completely mask the effect of attractive odors ^[26, 27]^. These studies utilized two odors each (ethyl acetate and benzaldehyde for fruit fly and isoamyl alcohol and 2-nonanone for *C. elegans*), however, the odors in these pairs trace their origins to various microbes or plants. Such laboratory-introduced combinations of odors, from disparate sources, may not have any ecological relevance for worms (or flies) and thus their combinations may not serve as food quality codes in animals. In our study, we used an attractant and a repellent from the same source (*P. aeruginosa*) and see no odor masking. Instead, we find strong evidence for worm’s ability to robustly track the precise ratio of attractant to repellent. Such a mechanism, to track ratios of repellent to attractant, has been recently reported in *D. melanogaster* as well ^[28]^.

Is the adaptable diet preference behavior of *C. elegans* reported here unique to *Pseudomonas*? We do not believe that to be the case. *Pseudomonas* and *Serratia* species are found in the natural habitat of *C. elegans*, so are other microbes that can produce a variety of attractive and repulsive cues. To identify other odor-based diet quality codes, a careful investigation of the odors generated by each microbe, followed by an evaluation of the worms’ response to odors, alone and in a mix, would be instructive. The ability to use specific odors from a bouquet as a food quality code could be a widely used strategy in the animal kingdom, allowing them to find better food ensuring the survival of the species.

## ACKNOWLEDGMENTS

Some *C. elegans* strains were provided by CGC which is funded by the NIH Office of Infrastructure Programs (P40 OD01440). We thank Dr. Radhika Buddidhathi for the synthesis of hexanoic acid, 2-ethyl methyl ester.

## AUTHOR CONTRIBUTIONS

S.H.C. and V.S. supervised the project. D.P., R.S., and V.S. designed the study and performed the analysis. D.P. and R.S. performed Choice, and Chemotaxis assays, and GC-MS/MS experiments for the analysis of bacterial volatile profile. D.P. performed the GCaMP imaging under the supervision of S.H.C. D.P., R.S. and V.S. wrote the manuscript.

## DATA AVAILABILITY

This study includes no data deposited in external repositories.

## METHODS

### Strains and growth media

*Caenorhabditis elegans* strains used in this study were maintained at 20°C on *E. coli* OP50 seeded Nematode Growth media (NGM) plates with streptomycin, as described previously ^[29]^. Unless otherwise specified, all experiments were performed at 25 °C. The N2 (Bristol) was utilised as the *C. elegans* wild-type strain in all experiments. Key resources table (Table S1) lists all the strains and chemicals used in this study.

### Food preference Assay

Food preference assay was performed on 90 mm slow killing (SK) media plates with 2% agar ^[30]^. Plates were air-dried for 70 mins and stored at room temperature for at least 48 h before using them. 25 µl of overnight grown bacterial cultures at 37°C (rotator) were spotted 1 cm away from the periphery on the diametrically opposite end of the plates (Figure 1A). These plates were then incubated at 37°C for 7 h. Before performing the assay, plates were kept at room temperature for the plates to cool down. The assay was performed using a synchronized population of adult worms. Worms were washed thrice with S-basal buffer and spotted at the centre of the plate. Scoring was done after 2 h. Preference index was calculated using the formula: *(Worms present on PA14 lawn - worms present on OP50 lawn)/Total worms*.

### Chemotaxis Assays

90 mm dishes of buffered agar were used to perform all the chemotaxis assays as described ^[14]^. Chloroform was used as the solvent control for all the odors used in this study. 2 µl of test or solvent were spotted on two opposite sides of the assay plate, (Figures 1C). Gravid worms were washed thrice with S-basal buffer and approximately 60-80 worms were spotted in the centre of the assay dish. 2 µl of 1M sodium azide was spotted near the test as well as solvent spot towards the periphery of the dish, as a paralytic agent to prevent switching of worms once a choice was made. The assay plates were incubated at 25°C for the duration of the experiment and the scoring was done after 2h as described previously ^[14]^. The chemotaxis index was calculated by the formula: *Chemotaxis Index (CI)= (Worms in test zone - Worms in control zone)/Total number of worms*. Each data point in these experiments represents one assay plate with 40-60 worms each.

For adaptation assays, synchronized population of adult worms were washed twice with S-basal buffer. Worms were spotted on an unseeded NGM dish with a minimal amount of buffer and air-dried for 5 minutes. This dish was placed on the top of another NGM dish with 4 spots of 3µL pyrrole each, or young lawn (8h), or old lawn (24 h) of PA14 required for adaptation. The two plates were sealed with parafilm and incubated at 25°C for 90 minutes. Worms were washed once with S-basal buffer before spotting them at the center of the chemotaxis assay dish.

### Calcium imaging

Calcium imaging in individual olfactory neurons expressing GCaMP (Table S1) was performed in a custom-designed microfluidic device ^[13]^. Transgenic worm expressing GCaMP family of genetically encoded calcium indicators in individual odor sensory neurons were trapped in the device and their calcium response was assayed under various dilutions of pyrrole. The dilutions of the chemicals were made using M9 buffer. GCaMP imaging was performed using a Zeiss inverted microscope using a Photometrics EMCCD camera. The imaging was performed by capturing stacks of TIFF files for 180 s at 10 frames sec^-1^ using Metamorph software. In a 180 s imaging session for each worm, the chemical stimulus was provided in the sequence-10 s stimulus OFF, 120 s stimulus ON followed by 50 s stimulus OFF state. The images were analyzed using MATLAB scripts to plot the change in fluorescence to the baseline F_o_ values. Bar diagrams were plotted as the average change in 10 s window after stimulus addition (time 11-20 s) and 10 s window after stimulus removal condition (time 131-140 s).

### Preparation of Hexanoic acid, 2-ethyl-, methyl ester

To a suspension of 10mL (6.26 mmol) of 2-ethyl hexanoic acid, 10 mL of methanol, 0.7 mL (9.72 mmol) of thionyl chloride were added. The reaction mixture was stirred at room temperature for about 12 h. The progress of the reaction was monitored periodically by testing the mixture by thin-layer chromatography (TLC). At the end of the reaction, the reaction mixture was concentrated, extracted into dichloromethane, and dried over anhydrous sodium sulphate to yield 0.6 grams (3.79 mmol). The identity and purity of Hexanoic acid, 2-ethyl-, methyl ester was confirmed using gas chromatography-mass spectrometry (GC-MS) analysis.

### SPME-GC-MS/MS analysis of volatiles produced by *P. aeruginosa*

An overnight culture of *P. aeruginosa* PA14 was used to seed with 7 spots of 50 µl spots on a 60 mm SK plate. The plates were then incubated for 8 h at 37°C. For the collection of volatiles, two *P. aeruginosa* PA14 spotted plates were sealed together using parafilm 1 hour prior to collection. The SPME fibre was inserted in between the 2 bacterial plates by puncturing a hole in parafilm between the plates with a needle as described previously ^[14]^. A solid-phase microextraction fibre (SPME; divinylbenzene/ carbon-WR/ polydimethyl siloxane, 80 µm; Agilent Technologies, Part no. 5191-5874) was used for the collection of volatiles. The fibre was exposed to the bacterial volatile for 30 minutes at room temperature. Immediately after collection, the SPME fibre was inserted into the GC injection port for desorption of the bacterial volatiles. The analysis was performed using 8890C gas chromatography (Agilent) interfaced with a 7000D GC/TQ. A capillary column HP-5MS ultra inert (30 m × 0.25 mm and 0.25 m, Agilent 19091S-433UI: 0245625H) was used for separation with Helium gas as the carrier gas at a constant flow rate of 1.5 ml min^−1^. The injector was kept in spitless mode. The column temperature program consisted of injection at 40°C, hold for 1 min, a temperature increase of 5°C min^−1^ to 170°C, followed by a temperature increase of 50°C min^−1^ to 270°C and hold for 2 minutes. The temperature of the inlet was 225°C. MS source and MS quadrupole temperature were maintained at 230°Cand 150°C, respectively. MS mass range used for the analyses was 30-300m/z. Volatiles were identified by comparing the mass spectra obtained with the mass spectral library of the GCMS data system, NIST 2017 version 2.3 (National Institute of Standards and Technology) mass spectral library.

### Statistical analysis

GraphPad Prism was used to perform all the statistical analysis. Either two-tailed unpaired *t*-test or one-way ANOVA, followed by Dunnett’s multiple comparison test was used to perform statistical analysis (mentioned in individual Figure legends). The annotations for significance according to P values is as mentioned – ns (not significant) P > 0.05; * P ≤ 0.05; ** P ≤ 0.01; *** P ≤ 0.001; **** P ≤ 0.0001. All the experiments have at least three replicates unless otherwise stated.

