## Supplementary Figures for "Pyrrole produced by *Pseudomonas aeruginosa* influences olfactory food choice of *Caenorhabditis elegans*"

### Supplementary Data

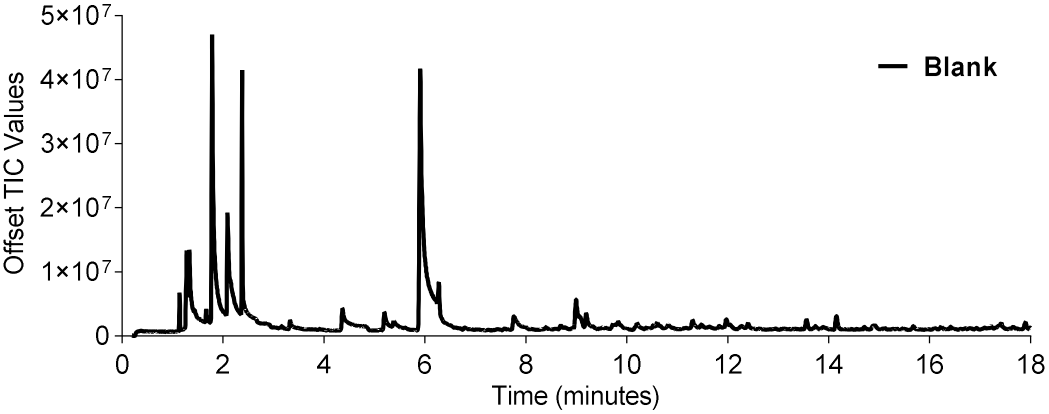


**Figure S1. GC-MS/MS analysis of volatiles from the media control dishes.**

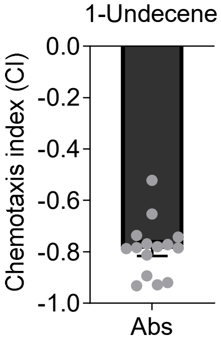


**Figure S2. 1-undecene produced by *P. aeruginosa* serves as a repellent for *C. elegans.***

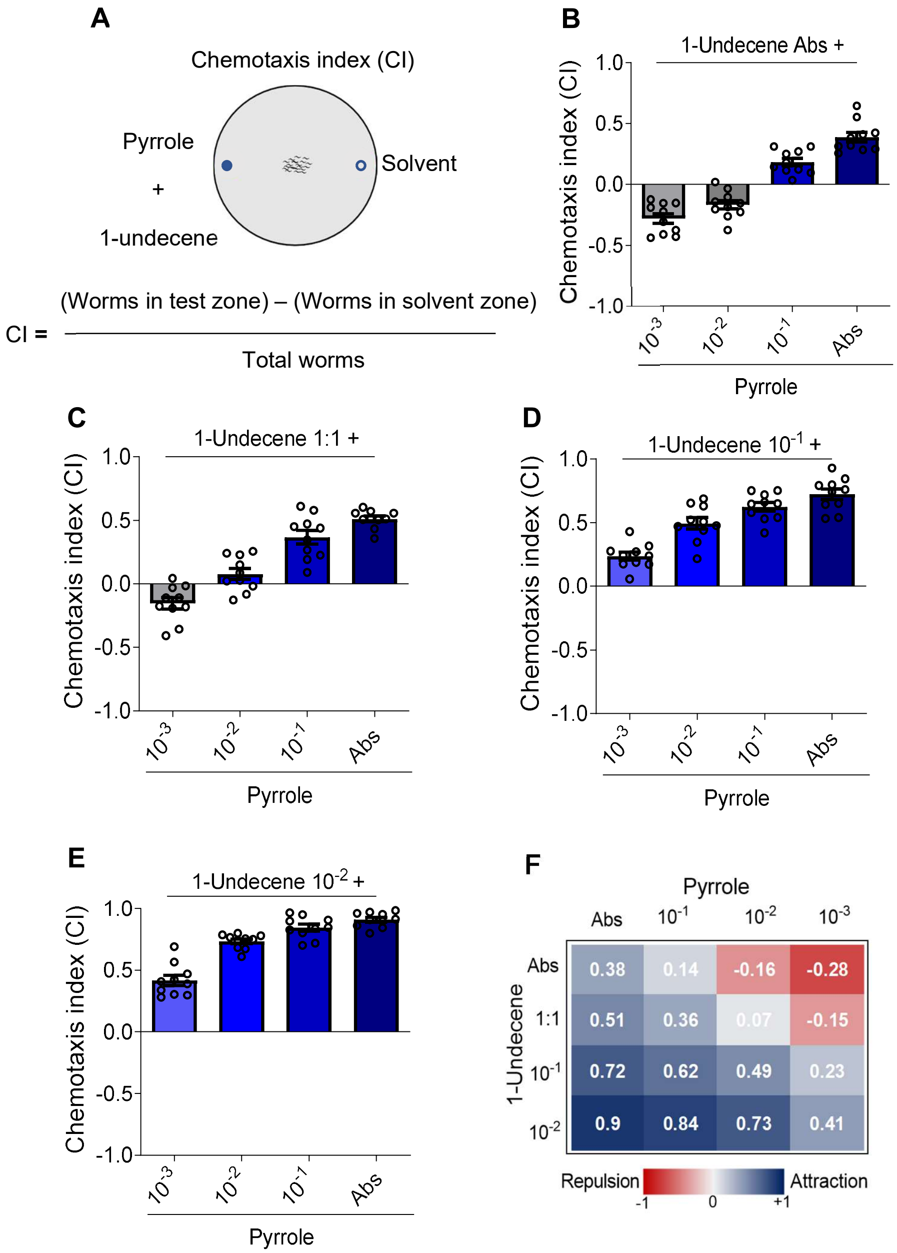


**Figure S3: Attractant to repellent ratio dictates chemotaxis response in worms.**

(A) Schematic for chemotaxis response in a complex olfactory landscape. (B-E) Chemotaxis response of N2 worms to a mixture of 1-undecene and varying concentrations of pyrrole against solvent control. Abs represents absolute concentration. n ≥ 3 assays, N ≥10 assay plates with 30-50 worms in each plate. (F) Heat map showing the average chemotaxis response of N2 worms to varying ratios of pyrrole and 1-undecene. n ≥ 3 assays, N ≥10 assay plates with 30-50 worms in each plate. **** P ≤ 0.0001 as determined by two-tailed unpaired *t*-test. Error bars indicate SEM.

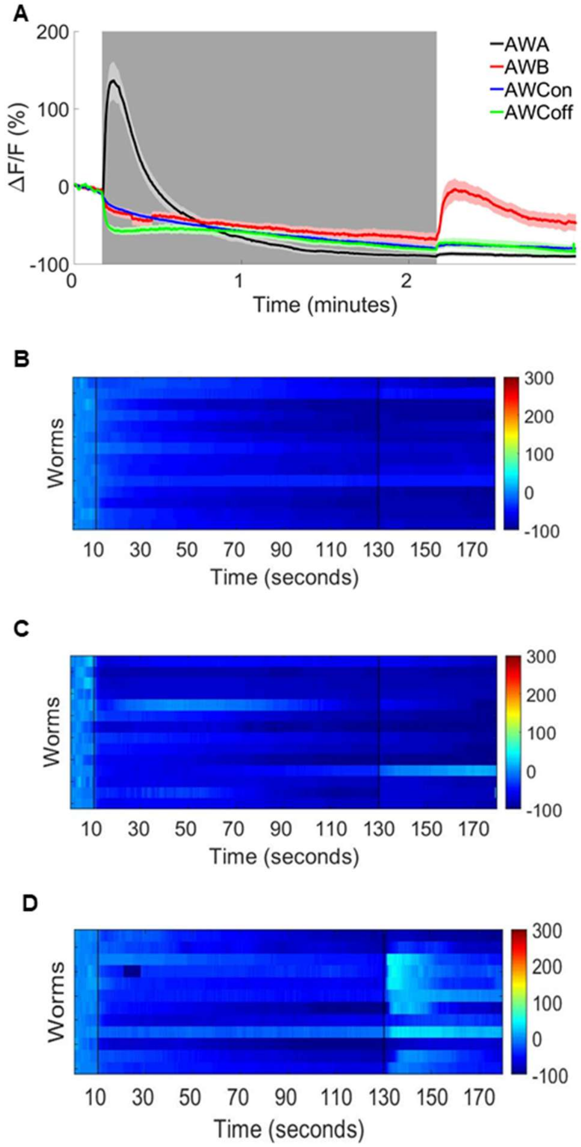


**Figure S4: Response of odor sensory neurons to the odor of pyrrole.**

(A) Average calcium responses of AWA::GCaMP2.2b, $AWC_{on}$:: GCaMP2.2b, $AWC_{off}$::GCaMP3 or AWB::GCaMP3 worms to pyrrole, recorded for 180 s, under stimulus between 11 s-130 s window shown in grey background. Heat map of calcium response in (B) $AWC_{on}$:: GCaMP2.2b, (C) $AWC_{off}$::GCaMP3 and (D) AWB::GCaMP3 worms. Each row represents an individual worm recorded for 180 s under 1:100 dilution of pyrrole, in 11 s-130 s window.

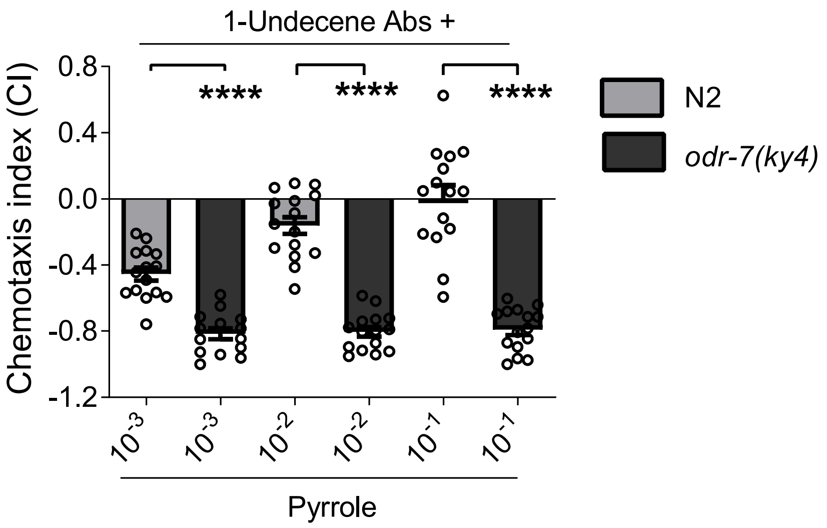


**Figure S5: Chemotaxis response of N2 worms and *odr-7(ky4)* mutant worms to mixture of 1-undecene and varying concentrations of pyrrole against solvent control.**

n ≥ 3 assays, N ≥10 assay plates with 30-50 worms in each plate. **** P ≤ 0.0001 as determined by two-tailed unpaired *t*-test. Error bars indicate SEM.

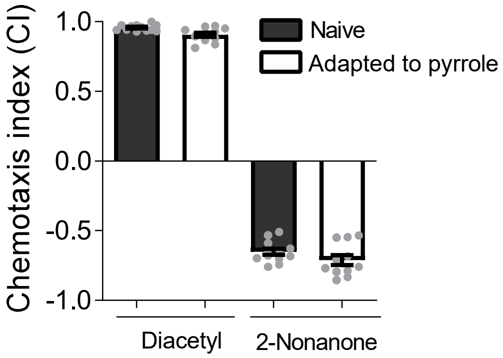


**Figure S6: Adaptation of *C. elegans* to pyrrole is highly specific.**

Chemotaxis response of naive worms and worms adapted to pyrrole to an attractive odor, diacetyl, and a repulsive odor, 1-nonanone.
